# Near-Zero Phase-Lag Hyperscanning in a Novel Wireless EEG System

**DOI:** 10.1101/2021.08.04.454932

**Authors:** Chun-Hsiang Chuang, Shao-Wei Lu, Yi-Ping Chao, Po-Hsun Peng, Hao-Che Hsu, Tzyy-Ping Jung

**Affiliations:** Research Center for Education and Mind Sciences, College of Education, National Tsing Hua University, Hsinchu, Taiwan; Institute of Information Systems and Applications, College of Electrical Engineering and Computer Science, National Tsing Hua University, Hsinchu, Taiwan; Artise Biomedical Co., Ltd., Hsinchu, Taiwan; Department of Biomedical engineering, National Yang Ming Chiao Tung University, Hsinchu, Taiwan; Graduate Institute of Biomedical Engineering, Chang Gung University, Taoyuan, Taiwan; Department of Computer Science and Engineering, National Taiwan Ocean University, Keelung, Taiwan; Institute for Neural Computation and Institute of Engineering in Medicine, University of California,San Diego, La Jolla, USA

**Keywords:** EEG, Hyperscanning, Brain-Computer Interface (BCI), RJ45, Radio Frequency (RF), Amplifier, Analog-to-Digital Converter (ADC), Timestamp, Sampling Clock, Brain Connectivity, Phase Locking Value, Signal Similarity

## Abstract

Hyperscanning is an emerging technology that concurrently scans the neural dynamics of multiple individuals to study interpersonal interactions. In particular, hyperscanning with wireless electroencephalography (EEG) is increasingly popular owing to its mobility and ability to decipher social interactions in natural settings at the millisecond scale. To align multiple EEG time series with sophisticated event markers in a single time domain, a precise and unified timestamp is required for stream synchronization. This study proposed a clock-synchronized method using a custom-made RJ45 cable to coordinate the sampling between wireless EEG amplifiers to prevent incorrect estimation of interbrain connectivity due to asynchronous sampling. In this method, analog-to-digital converters are driven by the same sampling clock. Additionally, two clock-synchronized amplifiers leverage additional RF channels to keep the counter of their receiving dongles updated, guaranteeing that binding event markers received by the dongle with the EEG time series have the correct timestamp. The results of two simulation experiments and one video gaming experiment revealed that the proposed method ensures synchronous sampling in a system with multiple EEG devices, achieving near-zero phase-lag and negligible amplitude difference between signals. According to all of the signal-similarity metrics, the suggested method is a promising option for wireless EEG hyperscanning and can be utilized to precisely assess the interbrain couplings underlying social-interaction behaviors.

## 1. Introduction

Brain–computer interfaces (BCIs) are considered a “next wave” technology [1]. The aim of the Neural Engineering System Design (NESD) program [2] of the Defense Advanced Research Projects Agency is to develop a high-resolution bidirectional BCI capable of providing precise and effective communication between humans and computers. In industry, technology companies [3] have launched ambitious programs and made progress in the development of BCI-assisted technologies, promoting the use of BCIs in real-world situations. Sensing and processing technology have advanced at a remarkable pace, and these advancements are influencing the future of BCIs. Incorporating social interaction [4, 5] into designs for deciphering brain mechanisms and mental processes in complicated natural situations is one of the emerging trends in real-world applications of BCIs. Such new BCIs, which aim to combine input from multiple users [6], necessitate a millisecond-scale concurrent neuroimaging approach and great portability for interactive scenarios.

Hyperscanning [7-10] is a new tool for concurrently exploring the brain functions of multiple people, and it’s been widely used to investigate the interbrain (de)synchronization that underpins social interactions [11]. Many joined-brain couplings have been revealed in situations where friends, strangers, colleagues, musicians, lovers [12], teachers and students, and mothers and children participating in interactive tasks such as singing [13], gaming [14], and video watching [15]. The hemodynamic or neuroelectric (de)synchrony between interacting brains is associated with coordinated behavior, shared cognition, and affective communication, as determined by functional magnetic resonance imaging (fMRI), functional near-infrared spectroscopy (fNIRs), magnetoencephalography (MEG), or electroencephalography (EEG). Among these regularly used neuroimaging tools, EEG provides temporal resolution and high mobility, making it the best tool for hyperscanning, particularly in a real-life setting.

EEG-based hyperscanning can conveniently read signals from multiple brains on the millisecond scale. The first relevant experiment was completed by Babiloni *et al*. [16], who recorded data from a group of participants wearing EEG headsets while playing cooperative games. EEG hyperscanning has become increasingly popular, but the synchronization method used in EEG hyperscanning has not yet been standardized. Several studies [9, 17, 18] have used an external trigger to synchronize the multiple data streams. Bækgaard *et al*. [17] employed spontaneous eye-blinking signatures to align EEG signals with eye-tracking patterns. Artoni *et al*. [19] delivered a digital input through a transistor– transistor logic (TTL) port to EEG and electromyography (EMG) devices simultaneously, discovering a misalignment of ±5 ms and jitter of 1.7 ms in a 10-min recording. Xue *et al*. [18] also used TTL pulses for synchronization when temporally aligning an EEG device with an eye tracker.

Recent studies have used a new data streaming framework named the Lab Streaming Layer (LSL)[20] to stream, synchronize, and collect multiple time series and event markers from different devices and software packages, including EEG, fNIR, eye-tracking, and motion-capture devices as well as stimulus-presentation software. The built-in time synchronization facility in LSL, which is similar to the Network Time Protocol, associates each sample with a timestamp to synchronize all recorded data at submillisecond accuracy on a local network of computers. For computers prepared for data acquisition, synchronization can be achieved by remapping the timestamps of data read from the local clock of computers in accordance with measurements of the momentary offsets between computers. One study [21] used LabRecorder [20], the main recording application of LSL, to synchronize and centralize the streams of two wireless EEG systems on a network while research participants performed a word-by-word interaction task. LSL-synchronized EEG systems were used in another study [22] to explore the mental workload of one pilot flying and another pilot monitoring during a simulated flight. With the support of cross-platform development and the need for multimodal data streaming, an increasing number of new devices feature an LSL plugin to enable two-way communication between devices with submillisecond timing precision. This synchronization approach can efficiently align multiple streams by combining the clock offsets with the timestamps of remotely collected samples. However, timing errors caused by offset correction estimation, multithreading, buffering, and wireless transmission are inevitable. Unfortunately, nonsimultaneous sampling due to asynchronous hardware remains a critical problem; this can result in amplitude differences and phase shifts, leading to incorrect estimates of interbrain connectivity.

As illustrated in Fig. 1A, a basic EEG device consists of electrodes, an amplifier, an analog-to-digital converter (ADC), and a digital signal processor (DSP). The ADC is a critical component for sampling EEG signals; it receives a clock signal from either a crystal or an oscillator, which means that analog signals can be sampled at specific discrete time points. Although modern ADCs have accurate clock sources with a wide range of frequencies, maintaining zero deviation from the true periodicity of a presumably periodic signal may still be challenging. For example, the Texas Instruments ADS1299 [23] has a frequency tolerance of ±50 ppm at 25°C and frequency stability of ±50 ppm over the operating temperature; that is, an overall stability and tolerance budget of 100 ppm can result in a potential frequency error of 0.01%. Suppose that two identical EEG recording units have frequency drifts of 0 and +100 ppm, respectively (Fig. 1B); the sampling rate difference between these two units could reach 0.1 Hz if the original clock signals of their ADCs have a frequency of 2.048 MHz. In the case of a 20-Hz sine wave (Fig. 1C), the difference between the phase signals recorded by these two ADCs would increase by 7.2° over a 10-s recording. Therefore, asynchronous clocks could lead to an incorrect estimate in connectivity analysis techniques that rely on measuring either amplitude or phase similarity, such as phase-locking value (PLV) [24, 25], phase-locking index (PLI) [26], coherence [27], imaginary coherence (ImgCoh) [28, 29], partial coherence [30], partial directed coherence [31], and phase–amplitude coupling [32].

**Figure 1.**
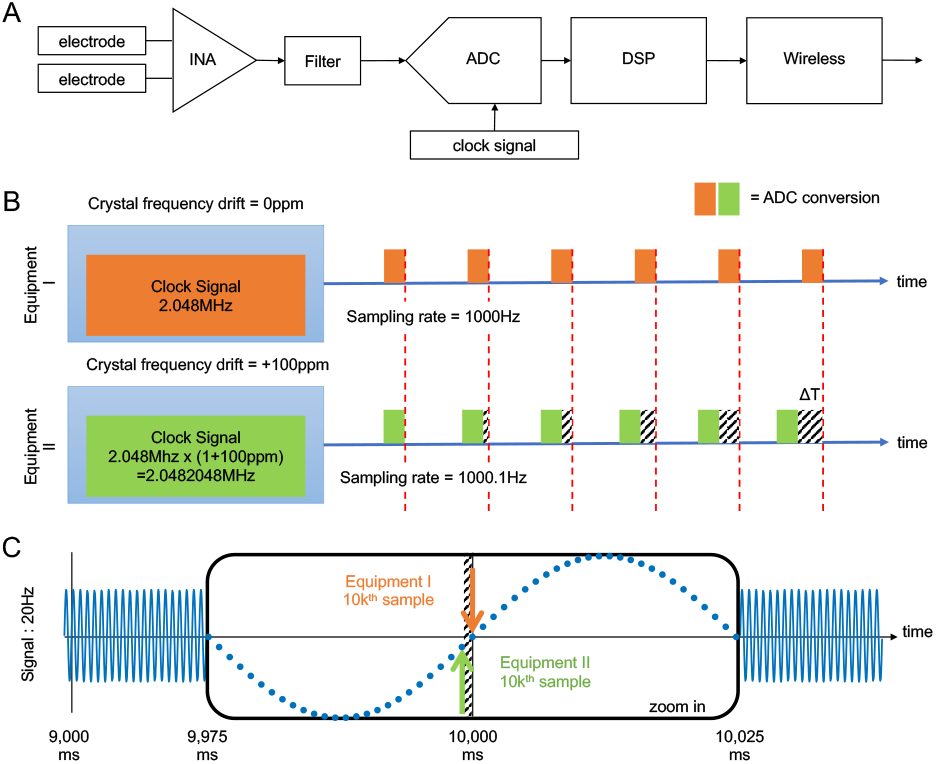
Asynchronous sampling. (A) Basic components of a wireless EEG device: electrodes, instrumentation amplifier (INA), filter, ADC, DSP, and wireless units. (B) Illustration of asynchronous sampling from two ADCs with a sample rate difference. Because the original clock signal has a frequency of 2.048 MHz, a crystal frequency drift of 100 ppm in Equipment II would cause a sampling error of 0.1 Hz. (C) Illustration of two ADCs sampling a 20-Hz sinusoidal signal. The two devices have sampling rates of 1000 and 1000.1 Hz, respectively; the enlargement shows that the 10 000th points sampled by these two devices are different.

In this study, we developed a new hyperscanning system capable of acquiring EEG signals wirelessly from multiple individuals while ensuring synchronous sampling across devices. Rather than simultaneously distributing an analog or digital synchronization signal between devices through a synchronization box [33] (Fig. 2A), the proposed method (Fig. 2B) coordinates the sampling across all EEG amplifiers by driving the ADCs with the same sampling clock via an RJ45 cable. Two simulation experiments and one video game experiment revealed that the proposed method could achieve zero-delay synchronization when measuring two identical signals. This study also compares the performance of several synchronization methods—the trigger-based, LSL-based, and proposed method—by calculating the amplitude difference and connectivity measures between the collected signals.

**Figure 2.**
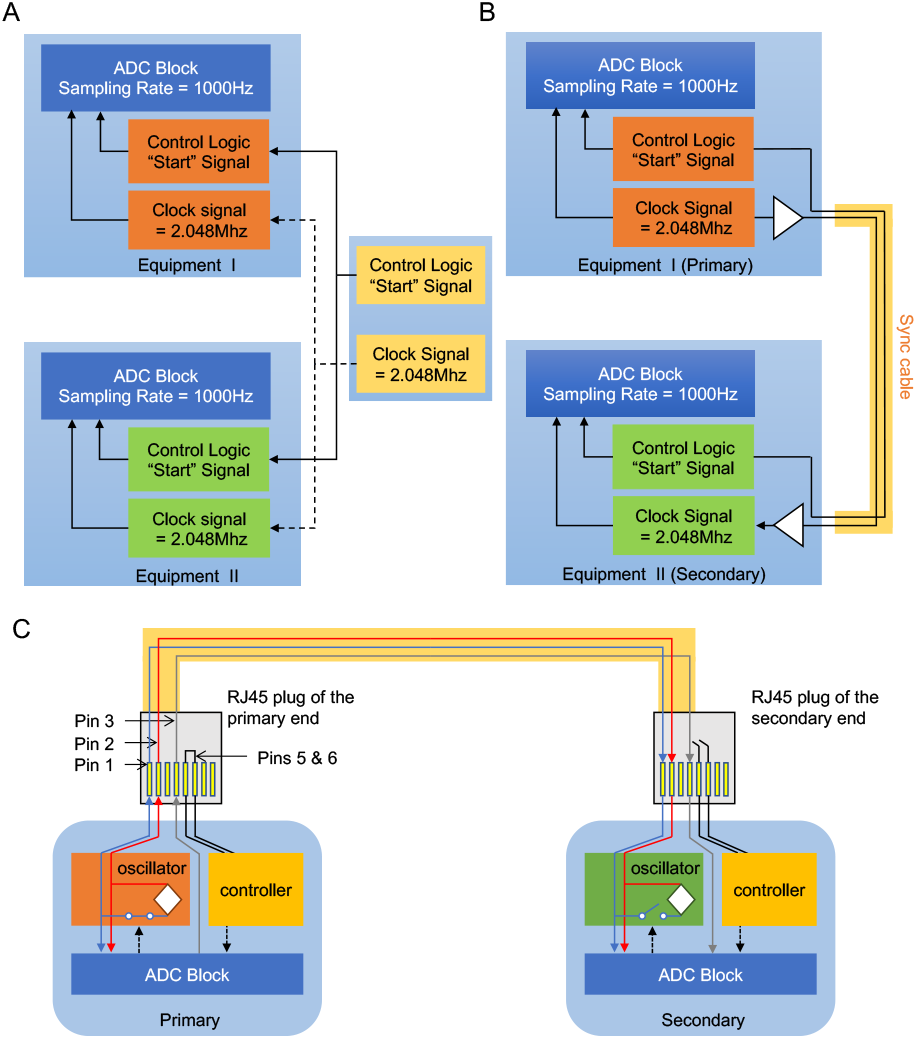
Proposed solutions for ADC synchronization. (A) An external control unit containing control logic and a clock signal generator that simultaneously sends a start signal and a clock signal to all ADCs. (B) Synchronization-cable design that enables devices to share the source of start and clock signals. One ADC, which coordinates the control and clock signals, is the primary device, and the additional ADCs are the secondary devices. (C) A custom-made RJ45 connector with an open-circuit/closed-circuit design for use as the synchronization cable between amplifiers. Pins 1 and 2 are designated for clock signal communication, Pin 4 is designated for start signal transmission, and Pins 5 and 6 are designated for switching between primary and secondary devices. Ideally, ADC conversions should occur simultaneously.

## 2. Methods

### 2.1 Proposed clock-synchronization method for hyperscanning

Fig. 2 shows two methods for synchronizing amplifier clocks to achieve hyperscanning. Instead of adding an external control unit to simultaneously send start and clock signals to all ADCs (Fig. 2A), this study proposed a synchronization-cable design that allows devices to share their source start and clock signals with each other. One of the amplifiers was designated as the primary device and coordinated the synchronization signal. The other amplifiers were the secondary devices and sampled the data following the pace set by the primary device.

This study employed a registered jack 45 (RJ45) cable, which is commonly used for connecting telecommunications and data equipment, to implement the proposed synchronization-cable design. As shown in Fig. 2C (left part), five of eight pins on the RJ45 cable were modified to transmit the synchronization signal and switch between the primary and secondary devices. Pin 4 was used to send the start-of-conversion signal from the ADC block of one device to another. Pins 1 and 2 were used to transmit clock signals between oscillators. Pins 5 and 6 were used to switch between the primary and secondary devices. Specifically, different designs of two J45 connector plugs were used to differentiate the primary and secondary devices. Pins 5 and 6 along with the controller formed a closed circuit at the “primary end,” generating a high voltage that signaled the oscillator to switch to the “primary mode.” In addition to continuing to transmit clock signals to the internal ADC, the oscillator was responsible for providing clock signals to external ADCs via Pins 1 and 2.

At the “secondary end,” as shown in Fig. 2C (right part), Pins 5 and 6 along with the controller formed an open circuit, generating a low voltage in the controller that signaled the oscillator to switch to the “secondary mode.” In this mode, the oscillator was unable to provide clock signals, and the ADCs received clock signals from the primary device via Pins 1 and 2 instead. The proposed design, utilizing a 3-m RJ45 fiber optic cable, would theoretically result in a propagation delay of approximately 50 ns, including 40 ns of delay caused by the drivers and receivers of the amplifiers.

### 2.2 Novel EEG system and timestamping

In this study, the proposed clock-synchronized hyperscanning method was implemented on a novel eight-channel wearable EEG device (Fig. 3). This EEG device comprised a dry/wet EEG electrode set (Fig. 3B), a wireless amplifier (Fig. 3C), and a receiving dongle (Fig. 3D). The amplifier was assembled using off-the-shelf chips and modules. With a powerful 64-MHz, 32-bit ARM Cortex M4 CPU, 1 MB of flash memory, and 256 kB of RAM, the host microprocessor and wireless communication featured a modular architecture that supported Bluetooth 5.0 and a proprietary 2.4-GHz RF. The customized RF receiver dongle was connected to a computer via a USB port, and the configured baud rate was set to 921 600 bits/sec to achieve high-speed transmission. Event trigger inputs were passed through an RS232 serial port on the receiver dongle for synchronization. A state-of-the-art chip provided by Texas Instruments (Dallas, TX, USA) guaranteed low input-referred noise and high 24-bit resolution for the delta-sigma ADC. Raw EEG data were sampled at 1000 Hz, streamed, and stored on a computer by using acquisition software developed using the Python language. The power source was a 500 mAh Li-polymer rechargeable battery, which provided sufficient power for 10 hours of continuous EEG recording.

**Figure 3.**
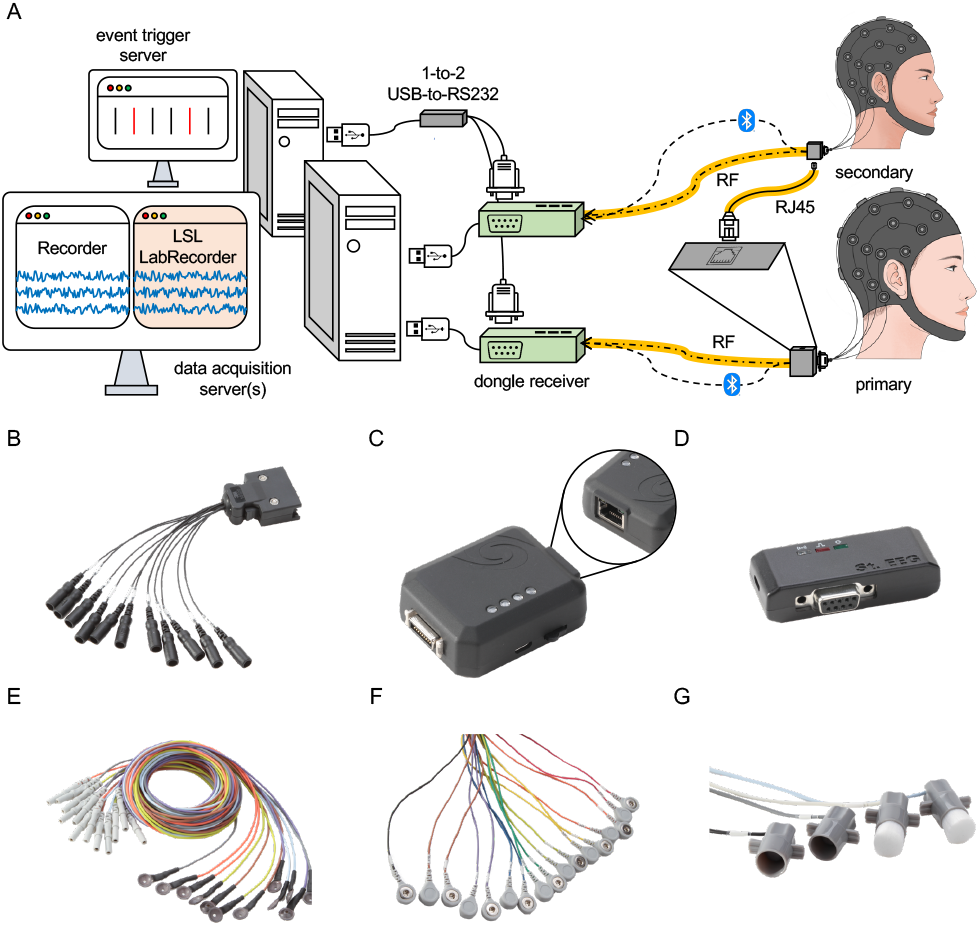
The proposed hyperscanning method implemented on two wireless EEG devices. (A) Hyperscanning setting: two amplifiers use the clock source shared by the primary amplifier. The clock data of amplifiers and EEG signals were transmitted to dongle receivers through RF and Bluetooth. This study proposed two approaches to unifying the data streams. In the first approach, two separate files are saved using the in-house acquisition software Recorder, and signals can be aligned in accordance with their timestamps. In the second approach, LSL is leveraged to stream signals, and LabRecorder is used to save them into a single file with a single time domain. (B) Electrode set: eight EEG electrodes, two references, and one ground channel. (C) an amplifier with an RJ45 socket for hyperscanning. (D) a dongle with an RS232 socket for receiving event markers. Three types of AgCl electrode are compatible with this wireless EEG system: (E) plate, (F) snap, and (G) sponge electrodes. These three electrodes can be used with conductive paste, disposable electrode pads, and saline water, respectively.

This study implemented an error-control mechanism, namely the Automatic Repeat Request (ARQ), into the wearable EEG hyperscanning system to enhance data transmission through the detection and retransmission of lost packets. Each packet was resent until it was successfully delivered or a time-out period had passed. Retransmission, even using the ARQ mechanism, might generate delays that might be misaligned with delays in other streams, such as an event marker. To address this problem, the proposed wireless transmission separated the recorded data into two streams. First, the EEG data stream included the timestamps were transmitted using Bluetooth over a 2.4-GHz RF to the receiving dongle. Second, an additional stream of timestamps defined by the clock signal was transmitted to the receiving dongle over another RF channel. As illustrated in Fig. 4, this additional RF transmission delivered timestamps every 100ms (10 Hz) over a 2.4-GHz frequency band. Two elapsed time counters in the transmitter (amplifier) and receiver (dongle) simultaneously started as the transmission began. Each timestamp packet updated the counter of the receiver, thus keeping the two counters synchronized. Even if a packet was lost during the RF transmission (i.e., if no timestamp was received by the dongle), the counter continued to work by providing accurate time information to the system. When the packet transmission resumed, the counter of the dongle was updated immediately. A timing error between counters would be extremely small unless severe signal interference or unknown RF transmission disruption was encountered.

In addition to the wireless transmission, two physical interfaces were established on the receiving dongle for data input/output. An RS232 interface was installed for accurately receiving event markers through the serial communication port. The proposed hyperscanning system used a 1-to-2 USB-to-serial RS232 adapter to enable event markers to be simultaneously received by two dongles (Figs. 3A and 4). Event markers were timestamped immediately once they arrived at the dongle to align them with the EEG signals. Because all system components shared common timestamps, the recording computer could integrate all streams into a single time domain.

**Figure 4.**
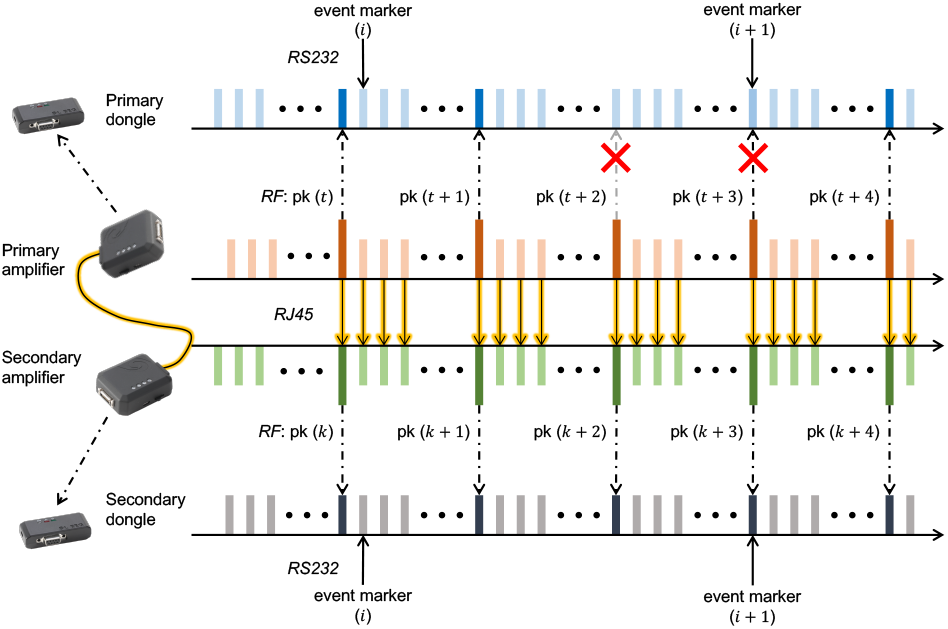
Timestamping of the proposed method. All components of the system share the same timestamps sourced from the primary amplifier. As soon as the recording starts, two clock-synchronized amplifiers send timestamp packets (denoted pk) through the RF channel to unify and update the counter of each dongle. This design binds event markers with the EEG time series in a unified time domain. Even if packet loss occurs during RF transmission, the built-in counter inside the dongle continues to oscillate to assure timing accuracy.

### 2.3 Comparison of synchronization methods

This study compared the stability and variability of signals acquired by different devices that were synchronized using the proposed approach versus other methods. As shown in Fig. 5A, the trigger-based method employed event triggers generated from an external device to mark time points on the acquired data. Specifically, a three-digit start event (code: 115) was communicated through PuTTY from a trigger PC through a 1-to-2 USB-to-RS232 converter to a recording PC with two recorders; then, this event was used to align signals for the subsequent analysis.

Fig. 5B illustrates the second synchronization method, in which the open-source LSL protocol was used for unified signal collection. The EEG recording software (EEG recorder) included LSL to receive signals from a wireless transmission module and used the LSL Stream Outlet function to broadcast these signals to a designated network. Suppose that a network has two available streams; the LSL LabRecorder [20] stores these streams along with their timestamps and clock offsets by reading them from individual local CPU clocks into a single .xdf file. Then, the MatLab function load_xdf.m is used to synchronize streams by correcting the timestamps, after which the amplitude differences and phase shifts between signals can be analyzed.

**Figure 5.**
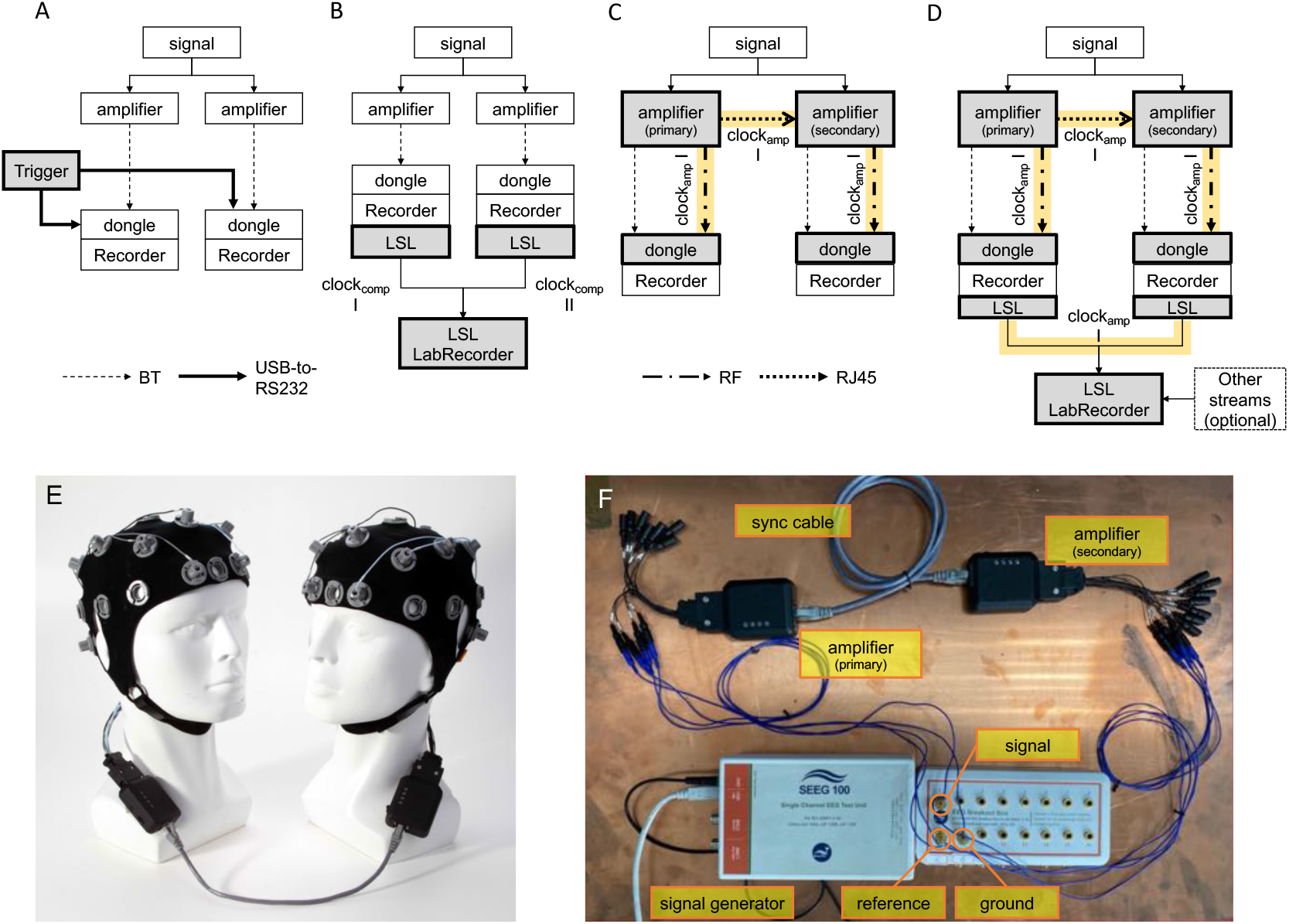
Synchronization methods for hyperscanning. The grey-highlighted areas represent the main steps of each technique for hyperscanning. (A) Trigger-based method: the signals collected by amplifiers are transmitted to their dongles via Bluetooth (dashed lines) and stored in separate files by using data acquisition software (EEG Recorder) along with the trigger. Then, the signals are aligned according to the trigger. (B) LSL-based method: the clock offset between two computers with EEG Recorder is estimated to remap the timestamps of each stream into a single time domain. Clockcomp I and Clockcomp II represent the CPU clocks of the computers. This synchronization method is completed by using the LSL Stream Outlet, LabRecorder, and load_xdf.m to send, receive, and merge data streams, respectively. (C) Proposed clock-synchronized method: the secondary amplifier adopts the clock source sent from the primary amplifier through an RJ45 cable. The RF channel is used to transmit the clock information from the primary amplifier, Clockamp I, to the dongle of the secondary amplifier, and this information is then used as the timestamp to align signals. The yellow areas indicate the information flow. (D) Proposed clock-synchronized method integrated with LSL for streaming: this design unifies all streams into a single time domain, where the amplifiers are still synchronously sampling. (E) Hyperscanning settings. (F) Hyperscanning setup for simulation experiments, in which simulated data generated using SEEG100 [34] were collected by each amplifier using one channel, two reference channels, and one ground channel.

The third method is the proposed approach (Fig. 5C), in which an RJ45 cable is used to synchronize all ADCs from the same clock source, allowing diverse EEG recording devices to be synchronized. The primary amplifier providing the clock source is designated the primary device. Another amplifier receiving the clock signal is designated the secondary device. These clock timestamps are then used to combine signals collected by different devices and stored by different recorders.

With the need for multimodal data streaming, this study further investigates if the synchronization performance would be maintained by combining the LSL with the proposed synchronization methods. Fig. 5D illustrates in this fourth method, which used an RJ45 cable synchronize the ADCs used for simultaneous sampling and timestamping and the LSL Stream Outlet function simultaneously to ensure that the signals were accessible on the designated network. By combining the advantages of hardware and software synchronization in an EEG recording system, the fourth method, if validated, could enable researchers to synchronize EEG streams with other modalities and facilitate a wide range of applications.

In summary, this study compares four synchronization methods: the trigger-based, LSL-based, clock-synchronized, and clock-synchronized with LSL methods; they used different timestamping approaches to synchronize the signals recorded from multiple devices and acquisition software packages.

## 3. Simulation results

### 3.1 Validation datasets

This study used two simulated datasets and one real EEG dataset to test the four synchronization strategies. The two simulated datasets comprised channels of 5-Hz and 30-Hz sine and square waves with 50 µVpp; which were continuously generated at a rate of 5000 pts/s using SEEG100 [34]. For the real EEG dataset, the replay function of SEEG100 was used to send signals to each amplifier. The test data were obtained from the EEG Motor Movement/Imagery Dataset (eegmmidb) [35] hosted on PhysioNet [36]. The validation dataset comprised FCz-channel signals from S001R04.edf, which had a sampling rate of 160 Hz and a dataset length of about 125 s (20 000 pts). The signal was replayed repeatedly for 30 min. To eliminate interference from the experimental environment and sources of variance between the amplifiers, the reference signal was averaged over two mastoid channels, and the ground signals collected in each amplifier were subtracted from the test signal. Each 30-min experiment was conducted five times to examine the robustness and time variability of the four synchronization methods.

A Butterworth notch filter was used to eliminate the 60-Hz and 120-Hz line noise, and a zero-phase filter was applied to the signals to further reduce noise in the signals. To test the synchronization performance, the absolute difference in amplitude between the signals recorded by the primary (p) and secondary (s) devices was calculated and averaged across all experiments; this difference is denoted 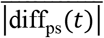 for each time *t*. The distribution of diff_ps_(*t*) was also obtained to determine the deviation of the differences from zero. Six signal similarity metrics were also obtained, which are typically employed to represent intra-or interbrain connectivity: the amplitude envelope correlation (EnveCorr) [37], power correlation (PowCorr) [38], circular correlation coefficient (CCorr) [39], PLV [25, 40], coherence (Coh) [38], and imaginary coherence (ImgCoh) [28, 29].

### 3.2 Similarity of signals recorded by different methods

To demonstrate the concept of asynchronous sampling illustrated in Figs. 1B and 1C, this study first compared the times series of simulated data recorded by amplifiers with and without clock synchronization. Fig. 6 shows 30-min 5-Hz sinusoidal and square waves collected by two amplifiers using the trigger-based method (Figs. 6A and 6C, respectively) and the proposed clock-synchronized method (Figs. 6B and 6D). When the trigger-based method was used, the two sinusoidal waves of the two devices appeared to be the same. However, at the start of the recording, there was a discrepancy in the time series (enlarged panel in Fig. 6A). This error escalated from ±5 to ±15 µV over the course of 30-min recording. The increasing error at the rising and falling edges of the square wave (Fig. 6C) indicated that the trigger-based method drifted from synchronous sampling with time. This result also suggested that that EEG amplifier manufacturing tolerances, which are tiny but not zero, could create significant variances when collecting millisecond-scale brain activity.

**Figure 6.**
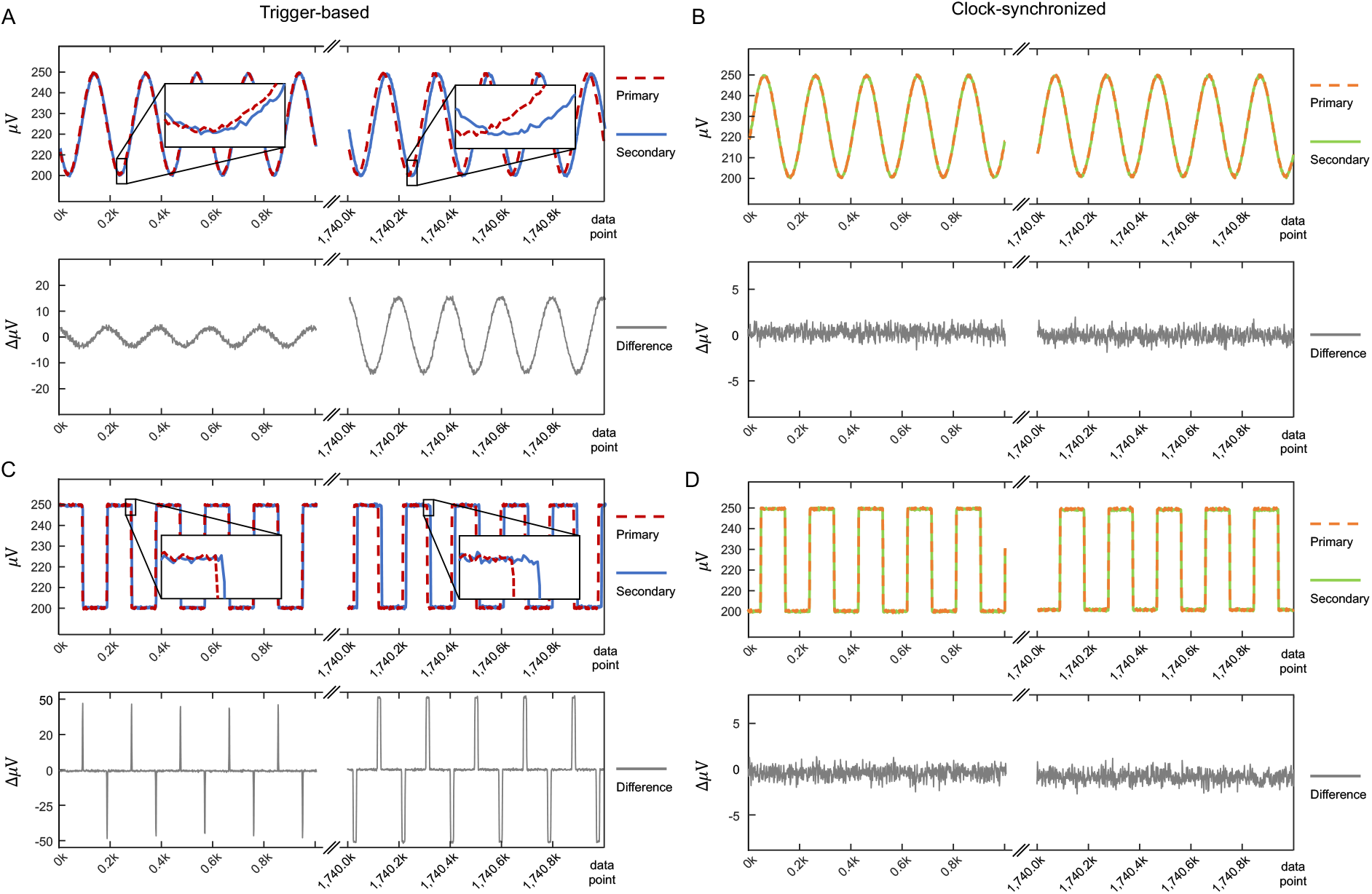
Sample 5-Hz sinusoidal and square signals collected by the two amplifiers through (A and C) the trigger-based and (B and D) the clock-synchronized methods. The upper panels show the amplitudes (µV) of two 30-min signals recorded by the primary and secondary amplifiers, and the lower panels show the amplitude differences (ΔµV) between them. A total of 1.8 million data points are included (1000 Hz ×60 s ×30 min).

By contrast, the proposed method, which used an RJ45 cable to synchronize amplifier clocks, achieved simultaneous sampling and flawlessly aligned signals (Figs. 6B and 6D) throughout the recording. The differences between signals remained minimal throughout the experiment. The robustness and stability of the proposed method were further supported by the following additional experiments.

Fig. 7 shows the differences between signals sampled at the primary and secondary devices when using the trigger-based, LSL-based, clock-synchronized, and clock-synchronized plus LSL method. Results for the 5-Hz simulated sinusoidal signals in Fig. 7A reveal that the 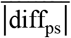 in the trigger-based method gradually grew to roughly 10 µV throughout the recording. When both amplifiers were turned on nearly simultaneously to record the signals and the signals were then aligned by using the starting trigger, 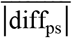 was minimized; this was also achieved in the system in which the two clocks were synchronized (blue and red traces). However, 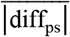 was only small at the beginning of he experiment. The signal difference between the sampling devices gradually increased as time passed. The amplitude difference between the two devices was ±10 μV towards the end of the recording. By contrast, when the other three synchronization methods were used, the amplitude differences of 5-Hz sinusoidal signals remained relatively remained reasonably minor and steady throughout the experiment. The diff_ps_ of the LSL-based and two clock-synchronized methods were ±5, ±2, and ±3 μV, respectively.

**Figure 7.**
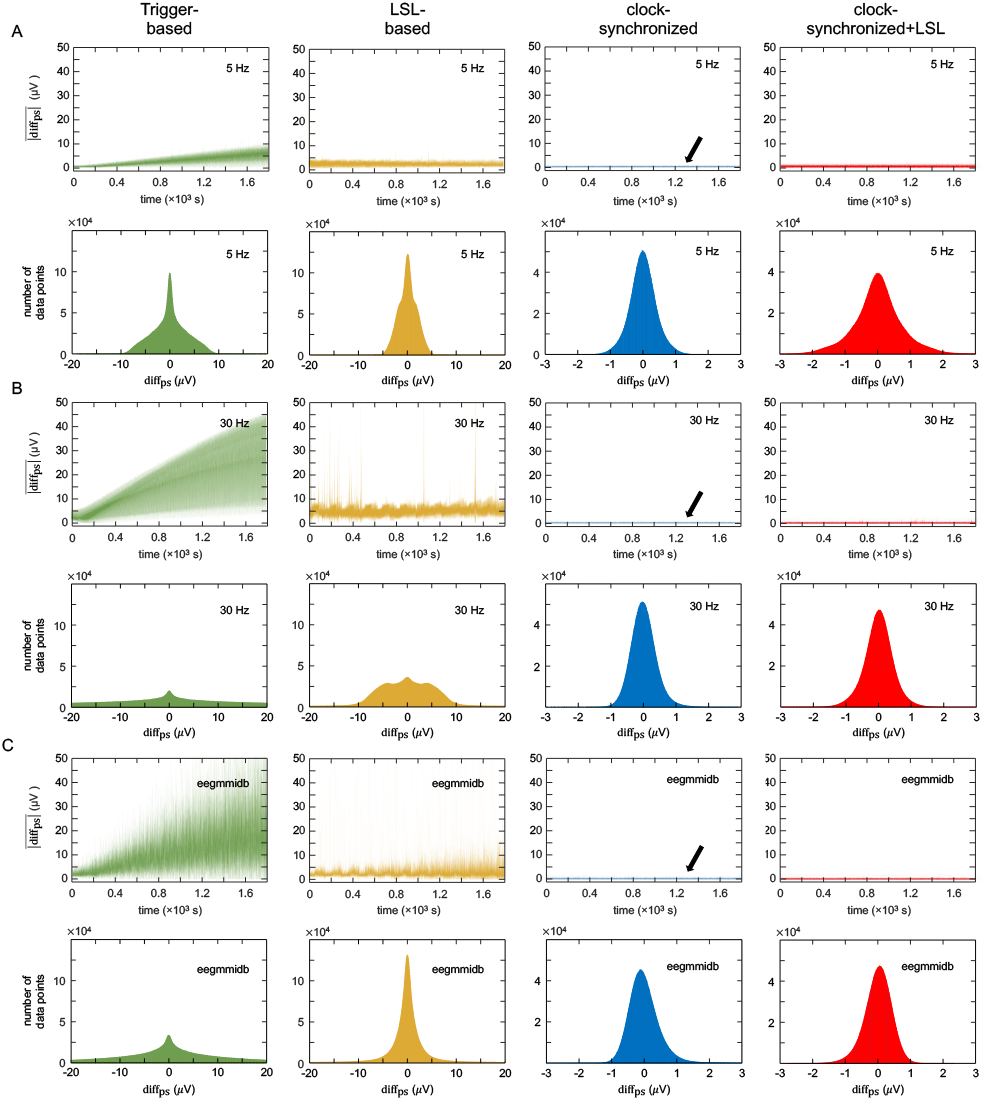
Amplitude differences (µ”V”) between two time series for (A) a 5-Hz sinusoidal siganls, (B) a 30-Hz sinusoidal siganls, and (C) eegmmidb. For each panel, subfigures, from left to right, show the results of hyperscanning using the trigger-based, LSL-based, and clock-synchronized methods. The upper panels show the absolute differences between signals averaged across five repeated experiments for each time point. The lower panels show histograms of the estimated differences between signals calculated for every time point and accumulated through five repeated experiments (1000 Hz ×; 60 s ×; 30 min ×; 5 times).

When the trigger-based method was used, the 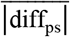 for the 30-Hz sinusoidal signals was up to approximately 45 μV, as shown in Fig. 7B. The diff_ps_ exhibited a platykurtic distribution, indicating considerable fluctuation in synchronization. When the LSL-based method was used, the 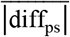 fluctuated within 10 μV throughout the recording, and most of the diff_ps_ values were distributed within ±10 μV. However, unaddressed issues such as asynchronous sampling and separate timestamping still resulted in significant differences between the data collected by the primary and secondary amplifiers. The proposed clock-synchronized methods sampled the 30-Hz signals perfectly, with the corresponding 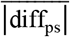 remaining constant throughout the recording, diff_ps_ distributed within ±1 μV.

Fig. 7C shows that the two amplifiers synchronized using either of the two proposed clock-synchronized methods could collect the real eegmmidb EEG signal [35]. The 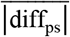 remained small and roughly constant, with the majority of the diff_ps_ values falling within ±1 μV. The least accurate approach was the trigger-based synchronization method, which had an initial error of 0 to 5 μV and an error that deteriorated as the recording progressed. The 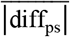 ranged from 0 to 50 μV, and the platykurtic distribution of diff_ps_ revealed inconstancies between the signals when using the trigger-based method. In the LSL-based method, the difference between the signals collected by the two amplifiers was smaller but could still be greater than ±10 μV.

Overall, the proposed methods outperformed the trigger-based and LSL-based methods in the hyperscanning of simulated sinusoidal signals and real EEG signals.

### 3.3 Synchrony between signals

In addition to measuring amplitude differences between signals, six commonly used connectivity measures— EnveCorr [37], PowCorr [38], CCorr [39], PLV [25, 40], Coh [38], and ImgCoh [28, 29]—were calculated to evaluate the performance of the four synchronization methods in terms of modeling the synchrony between signals recorded by different amplifiers. The estimated connectivity was benchmarked against the connectivity between two randomly generated signals and two identical signals. Two sets of 1 800 000 size-matched uniformly distributed random numbers ranging between 0 and 1 were generated to simulate two asynchronously recorded 30-min signals with a sampling rate of 1000 Hz. Moreover, the EEG signals from the previous validation experiment (Section 3.1) were duplicated to provide two identical and synchronous signals. As shown in Table 1, the EnveCorr, PowCorr, CCorr, PLV, Coh, and ImgCoh results for the random signals were 0.422, 0.443, 0.016, 0.288, 0.438, and 0.127, respectively. All connectivity measures of identical signals were 1.000, except for that of ImgCoh, which was 0.000.

**Table 1.**
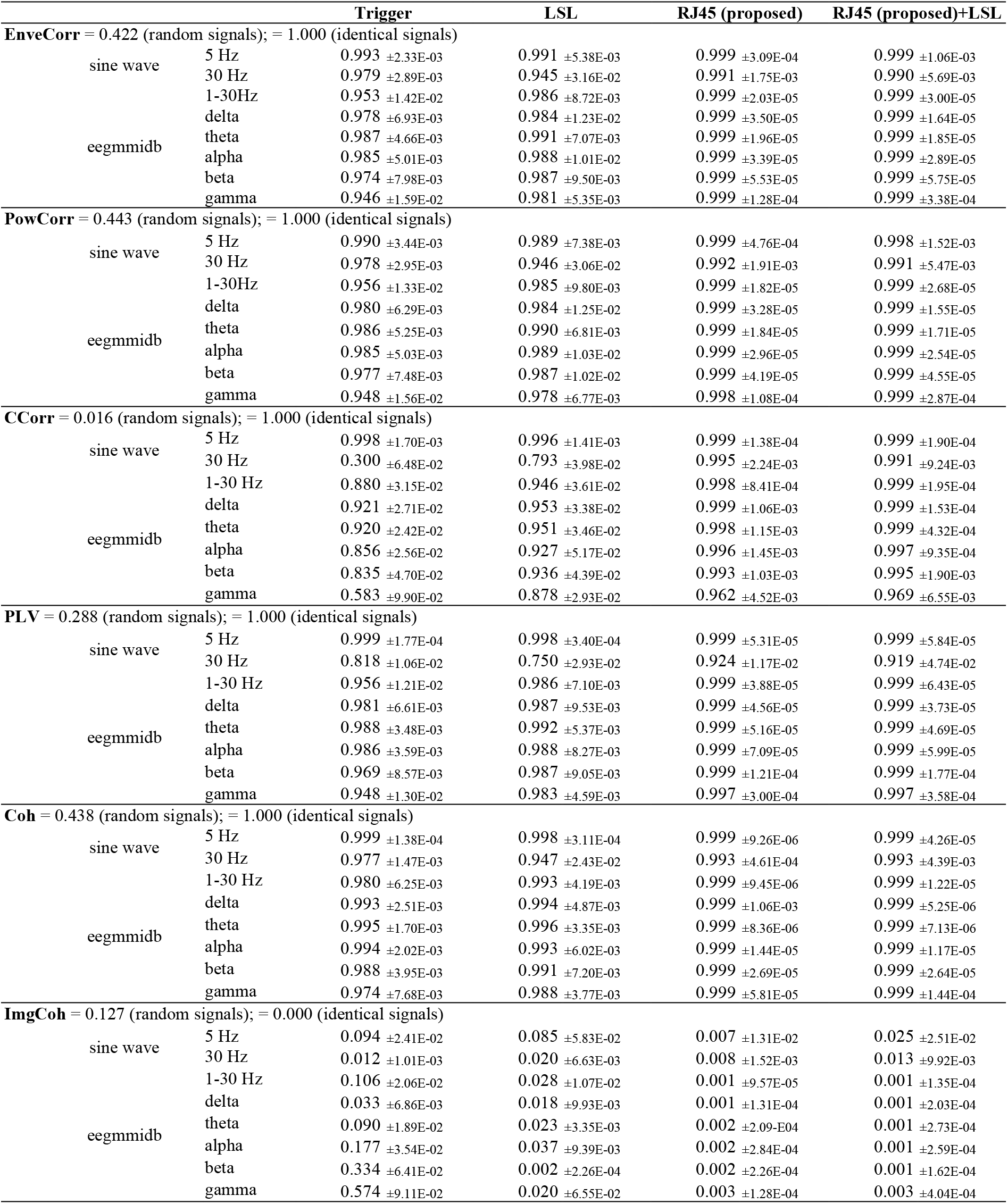
Connectivity between signals recorded using the four hyperscanning methods.

For the 5-Hz sine waves, the trigger- and LSL-based synchronization methods produced EveCorr values of 0.993 and 0.991, respectively, PowCorr results of 0.990 and 0.989, CCorr values of 0.998, and 0.996; and PLV results of 0.999 and 0.998. However, when recording high-frequency signals, as expected, the methods’ synchronization capability was substantially reduced, particularly for the 30-Hz sine wave and the gamma activity in eegmmidb. When the trigger-based method was used for the 30-Hz sinusoidal and eegmmidb EEG signals, EnveCorr ranged from 0.946 to 0.987, PowCorr from 0.948 to 0.986, CCorr from 0.300 to 0.921, PLV from 0.818 to 0.988, and Coh from 0.974 to 0.995. When the LSL-based method was employed for the 30-Hz sine wave and eegmmidb EEG signals, EnveCorr ranged from 0.945 to 0.991, PowCorr from 0.946 to 0.990, CCorr from 0.793 to 0.953, PLV from 0.750 to 0.992, and Coh from 0.947 to 0.996.

By contrast, when using the proposed hyperscanning methods, EnveCorr ranged from 0.990 to 0.999, PowCorr from 0.991 to 0.999, CCorr from 0.962 to 0.999 (except for the 30-Hz sine wave), PLV from 0.919 to 0.999, and Coh from 0.993 to 0.999 for the sinusoidal and eegmmidb EEG signals in different frequency bands. These results indicated that the signals recorded by the two amplifiers using the synchronized clocks were near identical.

Notably, the ImgCoh values of the identical signals and the signals collected by the proposed methods were approximately zero, suggesting that ImgCoh would fail to model the connectivity of two signals that were perfectly synchronous or had a zero phase-shift [41].

## 4. Experimental results

### 4.1 Pseudo-hyperscanning experiment

This study further used a computer card game called slapjack (heart attack) to investigate whether the proposed method could synchronize signals from multiple EEG amplifiers in a real experiment. As shown in Fig. 8A, this multiplayer game was developed using Unity3D and converted into a single-player mode. During the experiment, the card game was played using three modes, namely single player, cooperative, and competitive. The settings for the three modes were identical. During the game, a synthetic human voice was used to announce the name of a playing card (i.e., ace, two, three, …, jack, queen, or king) as a playing card was randomly drawn from a regular 52-card deck of cards and presented on the screen. The visual and auditory stimuli lasted for 1500 and 300 ms, respectively. The inter-trial interval was 0.5s. The subject was instructed to click a button as soon as possible if the displayed card matched the auditory stimuli. Two critical events, namely the stimulus onsets and response, were recorded. The reaction time was defined as the time between the two events. This event-related game requiring a quick sub-second response involved attention and inhibition skills.

**Figure 8.**
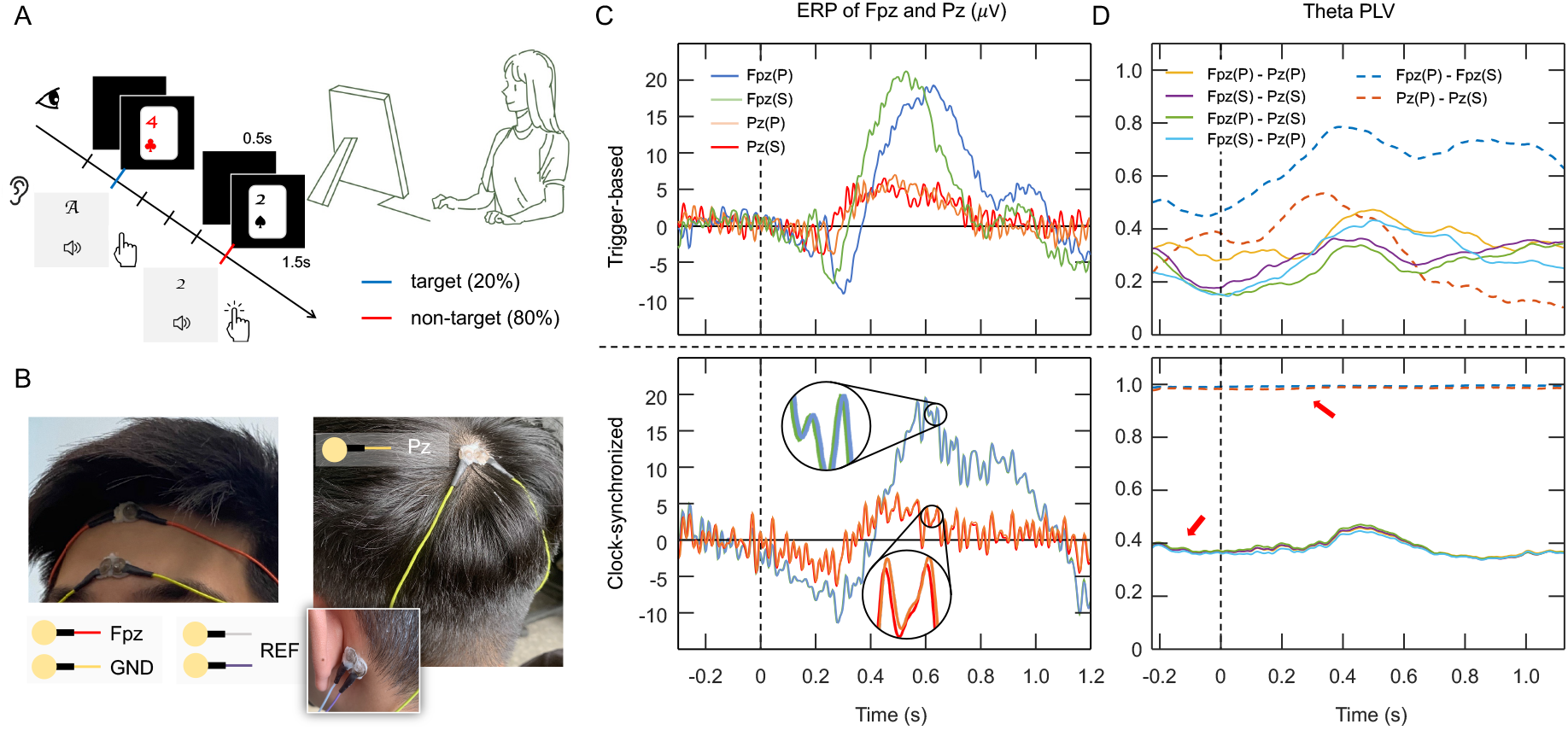
The hyperscanning validation experiment. (A) Experimental paradigm: the subject is instructed to click a button as soon as possible if the displayed card matches the auditory stimulus. The probability of the target was 20% (26 out of 130 trials), the duration of the target was 1.5 s, and the intertrial interval was 0.5 s. (B) For each amplifier, Fpz and Pz signals were collected with a ground (GND) electrode placed on the forehead and two reference (REF) electrodes placed on mastoids REF. Ample EEG conductive adhesive paste was applied to provide a bridge between connected electrodes, ensuring that the signals of neighboring channels were identical. (C) ERPs of Fpz and Pz recorded by the primary and secondary devices with the trigger-based method (upper panel) and the proposed method (lower panel). Each epoch was time-locked to the target event (vertical dashed lines) and segmented 0.3 s before and 1.2 s after the target. P and S represent the primary and secondary amplifiers, respectively. (D) Connectivity, evaluated using the event-related theta PLV, between two Fpz channels or two Pz channels. The two dashed traces indicate the PLVs between two connected channels. The four solid traces are the PLVs between pairs of remote channels. The upper and lower panels present the PLVs of the trigger-based and proposed methods. Each PLV was calculated over a sliding window of 150 pts (10% of the total data points).

During the game, a participant’s EEG activity was recorded using two separate wireless EEG devices (Fig. 5E). As shown in Fig. 8B, two EEG electrodes, one ground, and two references for each device were placed at the Fpz, Pz, forehead, right mastoid, and left mastoid, respectively. Ample EEG conductive adhesive paste was applied to bridge electrodes at these locations, enabling two devices to collect identical signals. This study then compared the event-related potentials (ERPs) of all four channels. Additionally, the event-related PLV was calculated to assess the signal similarity between all pairs of channels. The signals received by the two amplifiers were expected to be nearly identical. Therefore, the PLVs of cross-device connected channels [i.e., Fpz(P)/Fpz(S) and Pz(P)/Pz(S), where P and S represent the primary and secondary amplifiers, respectively] should have been approximately 1. In addition, the PLVs of any pair of non-connected channels [i.e., Fpz(P)/Pz(P), Fpz(P)/Pz(S), Fpz(S)/Pz(P), and Fpz(S)/Pz(S)] should have been identical. The synchronization performance of the trigger-based and proposed methods was compared.

The experimental protocol of this study was approved by the Research Ethics Committee at the National Tsing Hua University, Taiwan (NTHU-REC: 11001HT006). The participant gave his informed consent forms for inclusion before participating in the study.

### 4.2 ERP and PLV

The signals of connected sites should have been identical given the application of conductive paste (Fig. 8B). However, as revealed in Fig. 8C, the trigger-based method (upper panel), which used the target event to synchronize signals, failed to align the ERPs. The time delay between the signals recorded by the primary and secondary devices caused the PLVs to be underestimated. Fig. 8D indicates that the PLVs of Fpz(P)/Fpz(S) and Pz(P)/Pz(S) (two dashed traces on the upper panel) were far from 1. Consequently, the PLVs of the four Fpz/Pz pairs (two solid traces on the upper panel) evolved into different patterns, leading to misinterpretation of brain connectivity.

By contrast, the proposed method (lower panels of Figs. 8C and 8D) resulted in perfect synchrony when collecting EEG signals across all time points. The ERPs of the two Fpz channels and two Pz channels, which were recorded by two devices using the proposed clock-synchronized method, were nearly identical (lower panel of Fig. 8C). The PLVs of connected channels [i.e., Fpz(P)/Fpz(S) and Pz(P)/Pz(S)] were approximately 1 (two dashed traces on the lower panel of Fig. 8D). The PLVs of four Fpz/Pz pairs exhibited consistent event-related changes. Therefore, this near-zero phase-delay hyperscanning approach was able to assess valid coupling across brain regions.

## 5. Conclusion

The focus of BCI development has shifted in recent years from personalized use to group interaction. BCI communities should be able to grow thanks to hyperscanning and wireless technology. With the increasing research attention on social-interaction analysis, synchronization and hyperscanning have become crucial in new-generation BCIs. Synchronized EEG recording for multiple subjects demands the highest standard of time accuracy. When using wireless transmission, the latency of around 50–100 ms caused by Bluetooth is enough to cause the EEG analysis to be distorted. By sharing a clock source, this study used a hyperscanning method on two wireless EEG devices to accomplish synchronous sampling. A custom-made RJ45 cable connected the EEG amplifiers with an additional RF channel transmitting timestamps between the amplifier and its receiving dongle, enabling the EEG streams and event markers to be integrated into a unified time domain. The simulation and experimental results demonstrated the robustness and accuracy of the proposed method for collecting simulated and real EEG signals, and the results were superior to those achieved using the trigger- and LSL-based methods. Because the phase and amplitude discrepancies were so small, the suggested approach was able to record with minimal phase-lag and accurately show brain connectivity. Moreover, by applying different electrodes, such as ECG and EMG, the proposed system can simultaneously measure multiple physiological signals; this synchronous sampling technique can be utilized in a wide range of BCI applications.

## Acknowledgements

This work was supported by the Ministry of Science and Technology of Taiwan (project numbers: MOST 110-2636-E-007-018, 109-2636-E-007-022, 108-2321-B-038-005-MY2), and by the Research Center for Education and Mind Sciences, National Tsing Hua University. No funding source had involved in any of the research procedures.

